# A simple assay for inhibitors of mycobacterial oxidative phosphorylation

**DOI:** 10.1101/2023.08.08.552497

**Authors:** Serena A. Harden, Gautier M. Courbon, Yingke Liang, John L. Rubinstein

## Abstract

Oxidative phosphorylation, the combined activities of the electron transport chain (ETC) and adenosine triphosphate (ATP) synthase, has emerged as a valuable target for antibiotics to treat infection with *Mycobacterium tuberculosis* and related pathogens. In oxidative phosphorylation, the ETC establishes a transmembrane electrochemical proton gradient that powers ATP synthesis. Monitoring oxidative phosphorylation with luciferase-based detection of ATP synthesis or measurement of oxygen consumption can be technically challenging and expensive. These limitations reduce the utility of these methods for characterization of mycobacterial oxidative phosphorylation inhibitors. Here we show that fluorescence-based measurement of acidification of inverted membrane vesicles (IMVs) can detect and distinguish between inhibition of the ETC, inhibition of ATP synthase, and non-specific membrane uncoupling. In this assay, IMVs from *M. smegmatis* are acidified either through the activity of the ETC or ATP synthase, the latter modified genetically to allow it to serve as an ATP-driven proton pump. Acidification is monitored by fluorescence from 9-amino-6-chloro-2-methoxyacridine, which accumulates and quenches in acidified IMVs. Non-specific membrane uncouplers prevent both succinate- and ATP-driven IMV acidification. In contrast, the ETC Complex III_2_IV_2_ inhibitor telacebec (Q203) prevents succinate-driven acidification but not ATP-driven acidification and the ATP synthase inhibitor bedaquiline prevents ATP-driven acidification but not succinate-driven acidification. We use the assay to show that, as proposed previously, lansoprazole sulfide is an inhibitor of Complex III_2_IV_2_ while thioridazine uncouples the mycobacterial membrane non-specifically. Overall, the assay is simple, low cost, and scalable, which will make it useful for identifying and characterizing new mycobacterial oxidative phosphorylation inhibitors.

## Introduction

The genus *Mycobacterium* includes numerous pathogenic bacteria, the most notable of which being *Mycobacterium tuberculosis*, the causative agent of the disease tuberculosis (TB). Although mycobacteria can survive for an extended time in hypoxic conditions (Wayne and Lin, 1982), they are considered obligate aerobes. Oxygen serves as the final electron acceptor in oxidative phosphorylation, the combined activities of the electron transport chain (ETC) and adenosine triphosphate (ATP) synthase. The membrane-embedded protein complexes of the ETC couple oxidation of nutrients to the transport of protons across the mycobacterial inner membrane, producing an electrochemical proton motive force (PMF) that powers ATP synthase.

Unlike canonical ETCs found in mitochondria and many bacteria, the mycobacterial electron transport chain is highly branched (reviewed in Liang and Rubinstein, 2023). Nicotinamide adenine dinucleotide (NADH) is oxidized by at least two different NADH dehydrogenases: the proton pumping respiratory Complex I, as well as one or more non-proton pumping NDH-2 enzymes. Both Complex I and NDH-2 reduce menaquinone to menaquinol within the mycobacterial inner membrane. Two forms of respiratory Complex II (Sdh1 and Sdh2), and possibly fumarate reductase (Frd) functioning in reverse (Watanabe et al., 2011), serve as succinate:quinone oxidoreductases. A malate:quinone oxidoreductase (Mqo) can also contribute to the pool of reduced menaquinol in the membrane (Harold et al., 2022). Sdh1, Sdh2, Frd, and Mqo are not thought to contribute to the PMF directly, and Sdh2 may even harness the PMF to drive the endergonic reduction of menaquinone by succinate (Hards et al., 2020). Although mycobacteria can use electrons from menaquinol to reduce nitrate, fumarate, and hydrogen (Cook et al., 2014), oxygen is essential for mycobacterial growth (Wayne and Hayes, 1996; Wayne and Sohaskey, 2001). In the mycobacterial ETC, electrons are transferred from menaquinol to oxygen by two terminal oxidases: a supercomplex of respiratory Complexes III and IV (also known as cytochrome *bcc*-*aa*_3_ or CIII_2_CIV_2_) and cytochrome *bd*. The structures of CIII_2_CIV_2_ (Gong et al., 2018; Wiseman et al., 2018) and cytochrome *bd* (Safarian et al., 2021; Wang et al., 2021) suggest that they translocate four protons per electron and a single proton per electron, respectively.

The diarylquinoline drug bedaquiline was discovered in a phenotypic screen for compounds that inhibit growth of *Mycobacterium smegmatis* (Andries et al., 2005). Bedaquiline inhibits mycobacterial ATP synthase by binding with low affinity to the ring of c subunits in the enzyme’s rotor subcomplex (Andries et al., 2005; Preiss et al., 2015), with two high-affinity binding sites at the interface of subunit a with the c ring (Guo et al., 2021). Bedaquiline has become instrumental in the treatment of drug resistant TB (Cohen, 2017). Further, its discovery revealed that targeting oxidative phosphorylation can kill mycobacteria to treat TB. Subsequently, the second generation diarylquinolines TBAJ-876 and TBAJ-587 were developed, which exhibit improved binding to mycobacterial ATP synthase, reduced inhibition of the human Ether-à-go-go-Related Gene (hERG) channel, and decreased cLogP (Sutherland et al., 2019). These compounds are currently undergoing clinical trials. The imidazopyridine compound telacebec (also known as Q203) was discovered in a phenotypic screen of *M. tuberculosis* infected macrophages and found to inhibit CIII_2_CIV_2_ (Pethe et al., 2013). CIII_2_CIV_2_ activity can be replaced, at least in part, by cytochrome *bd* (Arora et al., 2014; Matsoso et al., 2005) and although treatment of *M. tuberculosis-*infected marmosets with a telacebec analog controlled disease progression and reduced lesion-associated inflammation it led to most lesions becoming cavitary (Beites et al., 2019). Nonetheless, treatment with telacebec decreased viable mycobacterial sputum load in humans in a phase 2 clinical trial (de Jager et al., 2020).

The activity of detergent-solubilized and purified complexes involved in oxidative phosphorylation can be measured in multiple ways. These methods include following the oxidation of substrates like NADH spectrophotometrically or monitoring reduction of oxygen with a Clark electrode. However, experiments with purified complexes are complicated by the tendency of soluble menaquinone analogues to autoxidize (Gong et al., 2018; Wiseman et al., 2018; Yanofsky et al., 2021; Zhou et al., 2021). Further, purifying ETC complexes is often time consuming and resource intensive, presenting a barrier to use of enzyme assays in the characterization of inhibitors of oxidative phosphorylation. Alternatively, oxidative phosphorylation can be measured with isolated mycobacterial membranes without purification of individual protein complexes. These membranes readily form inverted membrane vesicles (IMVs) upon resuspension in buffer that include intact ETCs capable of NADH-(Koul et al., 2007) or succinate-driven (Preiss et al., 2015) ATP synthesis. ATP synthesis can be followed with the enzyme luciferase, which uses ATP and oxygen to oxidize D-luciferin, producing oxyluciferin, AMP, carbon dioxide, and light, with the latter detected using a luminometer. However, even with IMVs, monitoring oxygen consumption is technically challenging and monitoring ATP synthesis requires either expensive reagents such as luciferase and luciferin (Koul et al., 2007) or complex procedures to denature proteins and quantify the remaining ATP (Haagsma et al., 2010). Further, distinguishing between inhibitors of ATP synthase and non-specific membrane uncouplers that dissipate the PMF to prevent ATP synthesis requires additional experiments. Despite these complications, ATP synthesis assays have been used by the company AstraZeneca to identify new mycobacterial ATP synthase inhibitors (Tantry et al., 2017).

In an alternative assay for mycobacterial ETC activity, IMVs can be added to a buffer containing the inexpensive fluorophores 9-amino-6-chloro-2-methoxyacridine (ACMA) or *N,N,N*′,*N*′-tetramethylacridine-3,6-diamine (acridine orange). Incubating the vesicles with an electron source, either succinate (Preiss et al., 2015), NADH (Koul et al., 2007), malate (Harold et al., 2022), or fumarate (Courbon et al., 2023) leads to the ETC pumping protons into the IMVs (Fig. 1A, *upper*). This acidification results in concentration of the fluorophore within the IMVs and quenching of its fluorescence, which can be restored by addition of an ionophore like nigericin to collapse the PMF. IMV acidification assays have been used extensively to demonstrate mycobacterial ETC activity (Courbon et al., 2023; Haagsma et al., 2010; Hards et al., 2015, 2018; Hotra et al., 2016), but in the absence of a complementary assay for ATP synthase activity they cannot identify ATP synthase inhibitors or distinguish ETC inhibitors from uncouplers.

**Figure 1.**
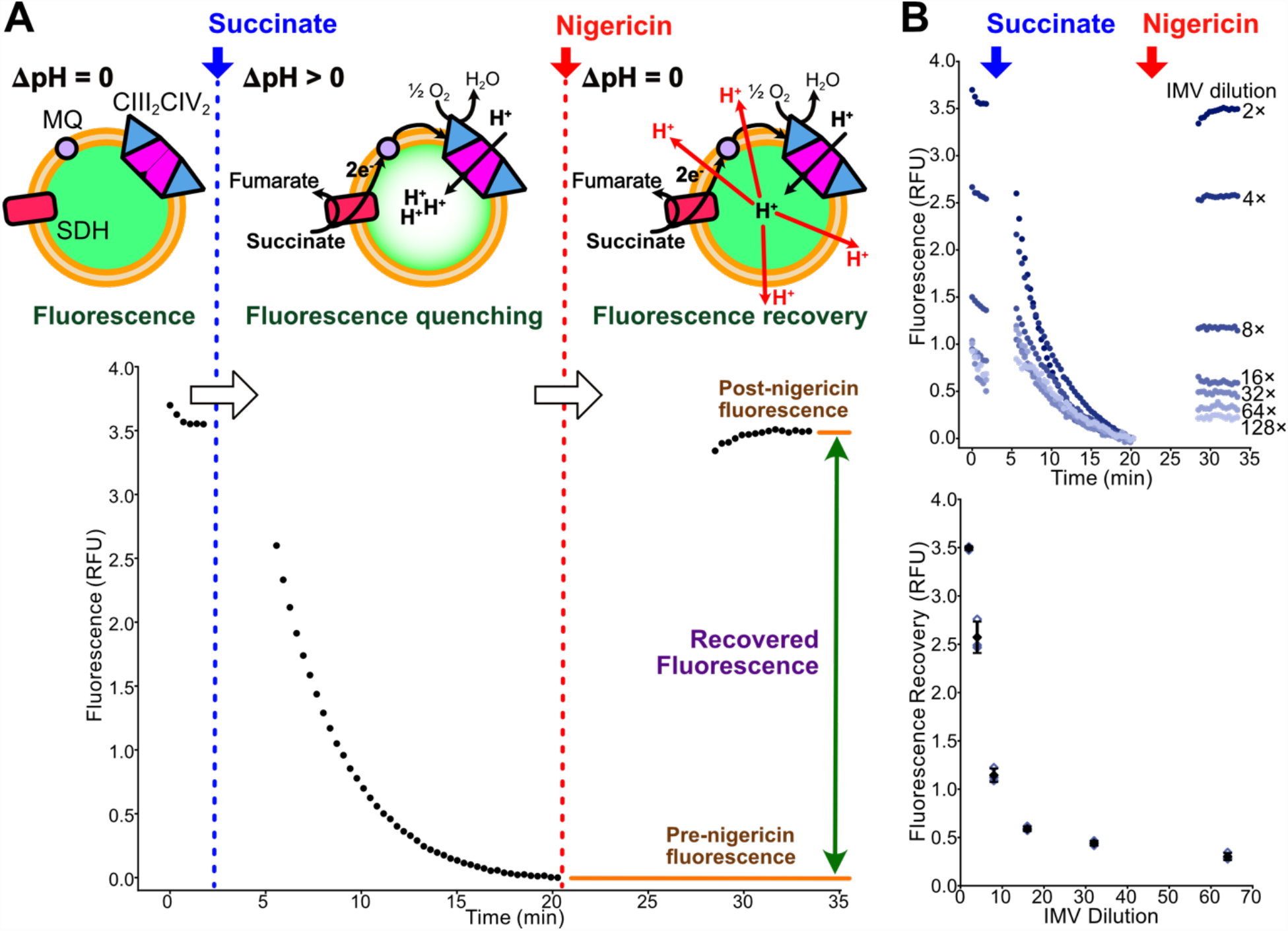
Quantification of succinate-driven acidification of *M. smegmatis* IMVs. **A**, Schematic of the IMV acidification assay with example data from succinate-driven acidification. SDH, succinate dehydrogenase; MQ, menaquinone. **B**, A dilution series (*upper*) and a plot of fluorescence recovery versus IMV dilution (*lower*) for succinate-driven acidification of IMVs shows decreased fluorescence recovery with more dilute samples. Open symbols show technical replicates. Filled symbols show the mean from n=3 technical replicates. Error bars indicate ± s.d. when shown.

Vesicles can also be acidified by ATP-driven proton pumps such as V-type ATPases (Bueler and Rubinstein, 2015) or mitochondrial ATP synthases working in reverse as ATPases (Groth and Walker, 1996). ATP hydrolysis by an ATP synthase can be detected by quantifying the release of free phosphate (Itaya and Ui, 1966; Taussky and Shorr, 1953) or with an assay that couples ATP hydrolysis to the oxidation of NADH, which can be monitored spectrophotometrically (Kornberg and Pricer, 1951). However, like many bacterial ATP synthases (Courbon and Rubinstein, 2022), mycobacterial ATP synthases are inhibited from working as ATP-powered proton pumps (Haagsma et al., 2010). In mycobacterial ATP synthase, ATP hydrolysis is inhibited by C-terminal extensions of the α subunits that prevent rotation of the rotor subcomplex in the hydrolysis direction (Guo et al., 2021). Modification of the genome of *M. smegmatis* to truncate the inhibitory extensions of the α subunits allows the enzyme to function as an ATPase (Guo et al., 2021) and chemical inhibition of ATP hydrolysis by the purified truncation mutant can be monitored with traditional ATPase assays (Courbon et al., 2023; Guo et al., 2021).

Here we show that ATP-driven acidification of IMVs from an *M. smegmatis* strain with truncated α subunits allows straightforward detection of mycobacterial ATP synthase inhibitors such as bedaquiline. Further, inhibitors of CIII_2_CIV_2_, such as telacebec, can be detected by their inhibition of succinate-driven IMV acidification, while non-specific uncouplers of the PMF prevent both succinate- and ATP-driven acidification. Surprisingly, NADH-driven acidification of IMVs is less sensitive to telacebec than succinate-driven acidification, even though both NADH and succinate should provide menaquinol to CIII_2_CIV_2_. As a result, the assay cannot distinguish succinate dehydrogenase inhibitors from CIII_2_CIV_2_ inhibitors. With the assay we show that lansoprazole sulfide (LPZS) blocks succinate-driven acidification of IMVs at sub-micromolar concentrations. LPZS is a metabolite of the gastric proton-pump inhibitor lansoprazole (LPZ) and was previously shown to inhibit CIII_2_CIV_2_ and have antimycobacterial activity (Rybniker et al., 2015). The assay also shows that thioridazine (THZ), a first generation antipsychotic drug that was found to inhibit mycobacterial NDH-2 (Rao et al., 2008; Weinstein et al., 2005; Yano et al., 2006), functions as an uncoupler of the PMF at micromolar concentrations. Owing to its simplicity and low cost, the assay should prove useful for identification and characterization of new inhibitors of mycobacterial oxidative phosphorylation.

## Results

### Fluorescence recovery allows quantification of IMV acidification activity

The absolute fluorescence measured from IMVs in a 96-well plate depends on fluorimeter settings and is sensitive to slight differences in the composition of the sample. Therefore, to investigate whether IMV acidification can be used to detect inhibition of oxidative phosphorylation, we first set out to find a way to compare fluorescence quenching between experiments. We did not attempt to quantify the pH within IMVs, but simply to measure the relative activity of the protein complexes that drive acidification. We found that this activity could be quantified most reproducibly by adding substrate, allowing ∼15 min for IMV acidification to quench fluorescence, and then measuring the recovery of fluorescence after adding a small volume of the potent H^+^/K^+^ antiporter nigericin to the sample (Fig. 1A, *green double headed arrow*). Using this quantification strategy and succinate-driven acidification, dilution of IMVs led to the expected decrease in fluorescence recovery (Fig. 1B, *upper*), although the fluorescence recovery was not linear with IMV concentration (Fig. 1B, *lower*). In our protocol for succinate-driven acidification, each sample in the 96-well plate contained 10 µL of IMVs prepared by resuspending membranes from 4 L of *M. smegmatis* culture in 10 mL diluted 2-fold before use. Consequently, with these conditions, a 4 L growth of bacteria provides enough material to perform up to 2,000 assays (or 500 assays/L of bacteria cultured).

### Succinate, NADH, fumarate, malate, and ATP can all drive IMV acidification

With the same preparation of vesicles used for succinate-driven acidification (Fig. 1B), we found that fumarate, malate, and NADH demonstrate the same concentration-dependent acidification of IMVs (Fig 2A-C). NADH:menaquinone oxidoreductase activity is likely the result of NDH-2 only, because the proton-pumping Complex I is almost undetectable in the conditions we used to culture *M. smegmatis* (Berney and Cook, 2010; Liang et al., 2023). Malate oxidation by Mqo also produces menaquinol (Harold et al., 2022), and as seen previously (Courbon et al., 2023) fumarate drives IMV acidification through a mechanism that is not completely clear but could involve a contaminating fumarase producing malate. Compared to succinate, addition of NADH, malate, and fumarate all result in faster and more extensive IMV acidification, as judged by the IMV dilution needed to produce a similar fluorescence recovery: 2-fold dilution for succinate, 4-fold for fumarate and malate, and 8-fold for NADH (Fig 1B and Fig. 2).

**Figure 2.**
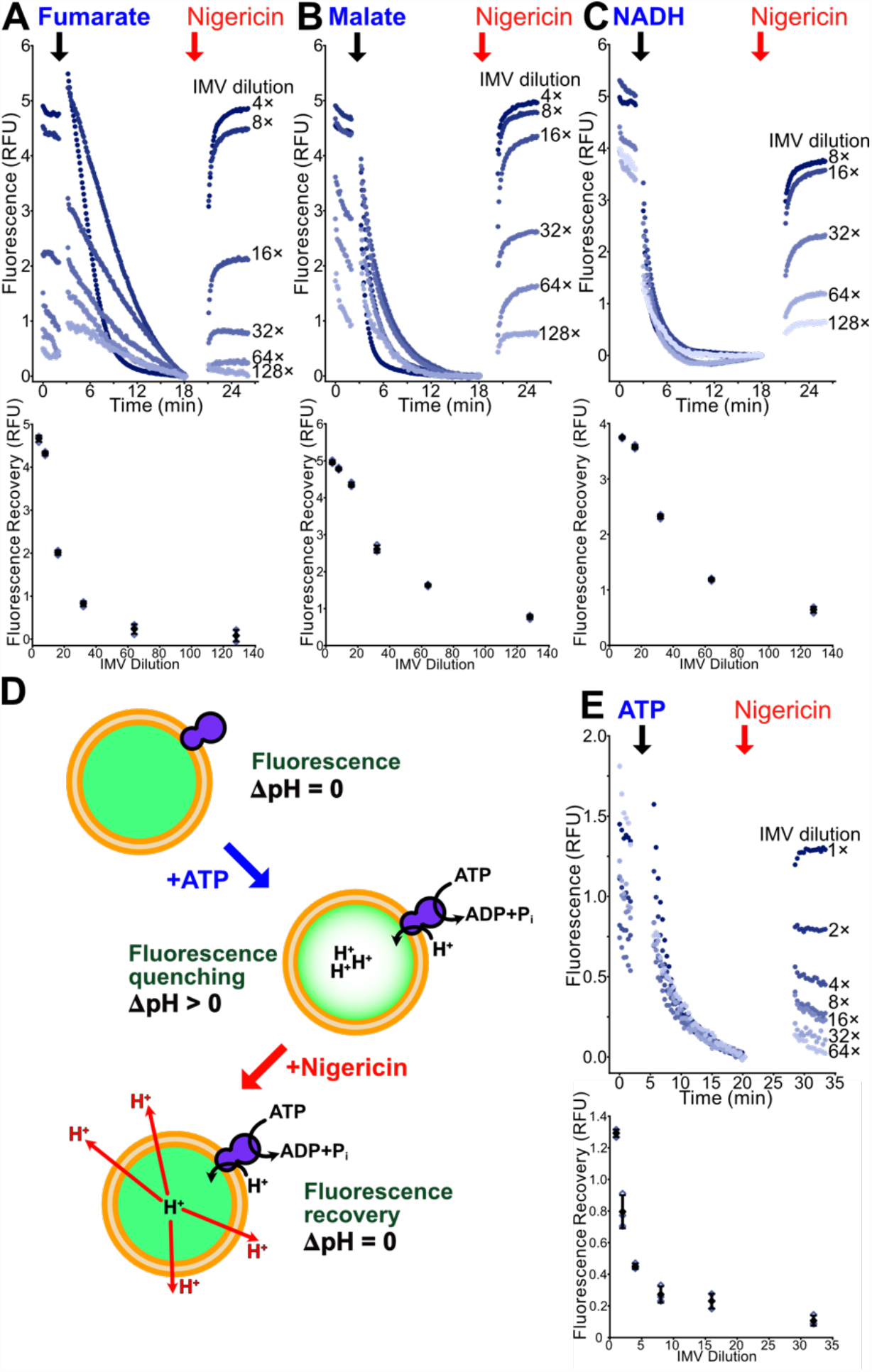
IMVs can be acidified with fumarate, malate, NADH, and ATP. **A**, Dilution series (*upper*) and a plot of fluorescence recovery versus IMV dilution (*lower*) for fumarate-driven acidification of IMVs. IMVs were prepared from the QcrB-3×FLAG *M. smegmatis* strain. **B**, Dilution series (*upper*) and a plot of fluorescence recovery versus IMV dilution (*lower*) for malate-driven acidification of IMVs. IMVs were prepared from the QcrB-3×FLAG *M. smegmatis* strain. **C**, Dilution series (*upper*) and a plot of fluorescence recovery versus IMV dilution (*lower*) for NADH-driven acidification of IMVs. IMVs were prepared from the QcrB-3×FLAG *M. smegmatis* strain. **D**, Schematic of the ATP-driven IMV acidification assay. **E**, Dilution series (*upper*) and a plot of fluorescence recovery versus IMV dilution (*lower*) for ATP-driven acidification of IMVs. IMVs were prepared from the *M. smegmatis* strain GMC_MSM2. Open symbols show technical replicates. Filled symbols show the mean from n=3 technical replicates. Error bars indicate ± s.d. when shown.

Next, we investigated whether ATP-driven acidification could be observed with IMVs prepared from an *M. smegmatis* strain where the α subunits of the ATP synthase had been truncated to allow ATP hydrolysis (Guo et al., 2021) (Fig. 2D). We found that this proton-pumping activity could indeed be observed (Fig. 2E). ATP-driven IMV acidification required using a higher concentration of IMVs in each well and was measured with 10 µL of IMVs prepared by resuspending membranes from a 3 L growth of *M. smegmatis* in 8 mL of buffer without further dilution. Although this condition requires more material per well than succinate-driven acidification, it still allows for ∼270 assays/L of bacteria cultured. This quantity of IMVs was used in all subsequent ATP-driven acidification assays. Like succinate, NADH, malate, and fumarate, fluorescence recovery from ATP-driven acidification also decreases as the IMVs are diluted.

### The CIII_2_CIV_2_ inhibitor telacebec inhibits succinate-driven IMV acidification

To determine if succinate-driven IMV acidification can be used to detect CIII_2_CIV_2_ inhibitors, we performed experiments with the well-characterized inhibitor telacebec (Q203) added to the sample at a range of concentrations. This experiment (Fig. 3A and B, with a replicate inhibitor dilution series in Fig. S1A) showed that telacebec can block succinate-driven IMV acidification, yielding an IC_50_ of ∼140 nM. This IC_50_ is comparable to the IC_50_ of 53 nM reported previously from measurement of purified CIII_2_CIV_2_ activity with an oxygen sensitive electrode (Yanofsky 2021). The near-complete inhibition of acidification by telacebec indicates that most of the acidification results from CIII_2_CIV_2_ activity rather than cytochrome *bd* activity. The same concentrations of telacebec did not inhibit ATP-driven acidification of IMVs (Fig. 3C), as expected from the compound’s specific inhibition of CIII_2_CIV_2_ rather than ATP synthase. Telacebec’s inhibition of succinate-driven acidification but not ATP-driven acidification also confirms that its effect does not arise from uncoupling of the PMF.

**Figure 3.**
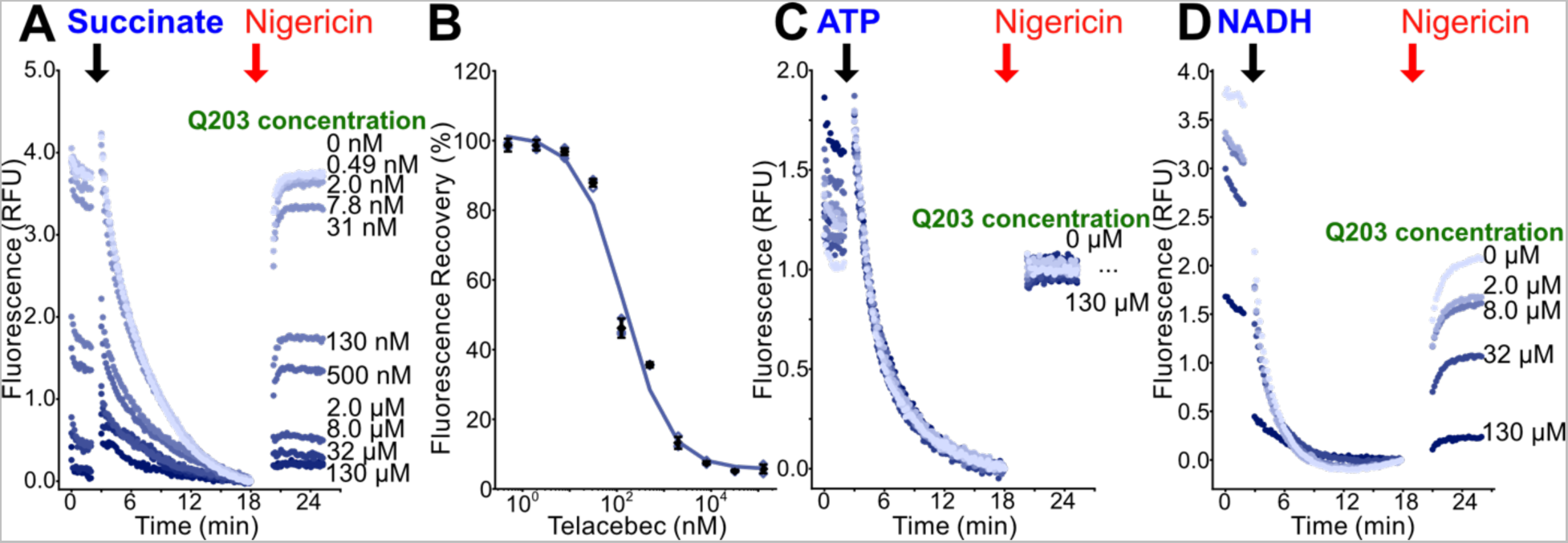
Telacebec inhibition of CIII_2_CIV_2_ can be detected by its effect on succinate-driven IMV acidification. **A**, Telacebec (Q203) at different concentrations inhibits succinate-driven IMV acidification. **B**, A plot of fluorescence recovery versus inhibitor concentration indicates an IC_50_ of ∼140 nM. **C**, Telacebec does not inhibit ATP-driven acidification of IMVs, showing that it is does not uncouple the PMF. **D**, Telacebec inhibits NADH-driven acidification of IMVs only at high concentrations. Open symbols show technical replicates. Filled symbols show the mean from n=3 technical replicates. Error bars indicate ± s.d. when shown.

Surprisingly, while NADH drives robust acidification of IMVs (Fig. 2C), this acidification could only be inhibited completely with high concentrations of telacebec and with IMVs diluted 16-fold relative to the concentration used for the succinate experiments. As described above, in the growth conditions for the bacteria used to prepare these IMVs NADH:menaquinone oxidoreductase activity is due almost entirely to the non-proton pumping NDH-2 (Berney and Cook, 2010; Liang et al., 2023). Whether menaquinol is produced by succinate dehydrogenase or NDH-2 it should contribute to the PMF in the same way via CIII_2_CIV_2_ and to a lesser extent cytochrome *bd*. Therefore, it is not clear why telacebec can block succinate-driven acidification almost completely, while only slightly inhibiting NADH-driven acidification. Unfortunately, the inability to detect inhibition of NADH-driven IMV acidification means that the assay cannot distinguish succinate dehydrogenase inhibitors from CIII_2_CIV_2_ inhibitors, although other assays exist for that purpose (Gong et al., 2018; Kun and Abood, 1949; Wiseman et al., 2018; Yanofsky et al., 2021).

### The ATP synthase inhibitor bedaquiline inhibits ATP-driven IMV acidification

We next tested whether IMV acidification could be used to detect inhibition of mycobacterial ATP synthase. Using IMVs with hydrolytically competent ATP synthase, we tested a range of concentrations of bedaquiline for inhibition of ATP-driven IMV acidification (Fig. 4A and B, with a replicate inhibitor dilution series in Fig. S1B). This analysis provided an IC_50_ of ∼34 nM, which is somewhat higher than the reported IC_50_ of ∼2.5 nM in an ATP synthesis assay with *M. smegmatis* IMVs (Koul et al. 2007), or the nanomolar inhibition of ATP hydrolysis by purified hydrolytically competent *M. smegmatis* ATP synthase (Guo et al. 2021). The higher IC_50_ values in the IMV acidification assay suggest that it is not as sensitive as the other assays, but it is still capable of easily detecting an ATP synthase inhibitor like bedaquiline. Bedaquiline did not show inhibition of succinate-driven IMV acidification, consistent with the compound being a specific inhibitor of mycobacterial ATP synthase that does not inhibit mycobacterial CIII_2_CIV_2_ (Fig. 4C). The lack of inhibition of succinate-driven IMV acidification also demonstrates the inhibition of the ATP-driven acidification is not the result of uncoupling the PMF.

**Figure 4.**
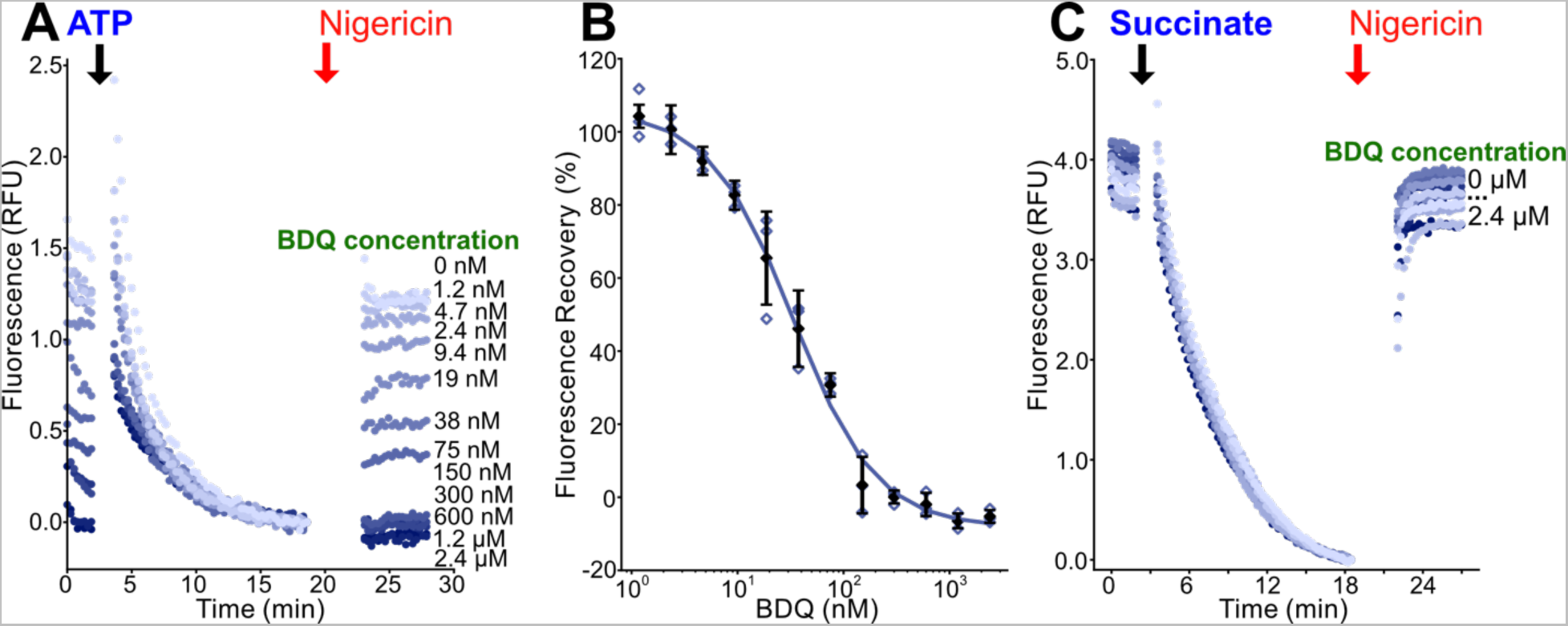
Bedaquiline inhibition of ATP synthase can be detected by its effect on ATP-driven IMV acidification. **A**, Bedaquiline (BDQ) at different concentrations inhibits ATP-driven acidification of IMVs. **B**, A plot of fluorescence recovery versus inhibitor concentration indicates an IC_50_ of ∼30 nM. **C**, Bedaquiline does not inhibit succinate-driven acidification of IMVs, showing that its effect on ATP-driven acidification is not a result of uncoupling the PMF. Open symbols show technical replicates. Open symbols show technical replicates. Filled symbols show the mean from n=3 technical replicates. Error bars indicate ± s.d. when shown.

### LNZS inhibits succinate-driven IMV acidification while THZ uncouples the PMF

Lansoprazole (LNZ; 2-{[3-methyl-4-(2,2,2-trifluoroethoxy)pyridin-2-yl]methanesulfinyl}-1H-1,3-benzodiazole) is a H^+^/K^+^-ATPase gastric proton pump inhibitor sold under the brand name Prevacid and used extensively in medicine. A metabolic product of lansoprazole, lansoprazole sulfide (LNZS), was found to have antimycobacterial activity owing to inhibition of CIII_2_CIV_2_ (Mdanda et al., 2017; Rybniker et al., 2015). LNZS showed inhibition of succinate-driven IMV acidification with an IC_50_ of ∼860 nM (Fig. 5A and B, with a replicate inhibitor dilution series in Fig. S1C). The compound did not inhibit ATP-driven acidification (Fig. 5C), indicating that inhibition succinate-driven acidification is the result of blocking electron transport chain activity rather than uncoupling the PMF. In contrast to LNZS, LNZ did not show inhibition of succinate-driven acidification (Fig. 5D) supporting the finding that LNZS, and not LNZ, is the active compound in inhibition of mycobacterial respiration (Rybniker et al., 2015).

**Figure 5.**
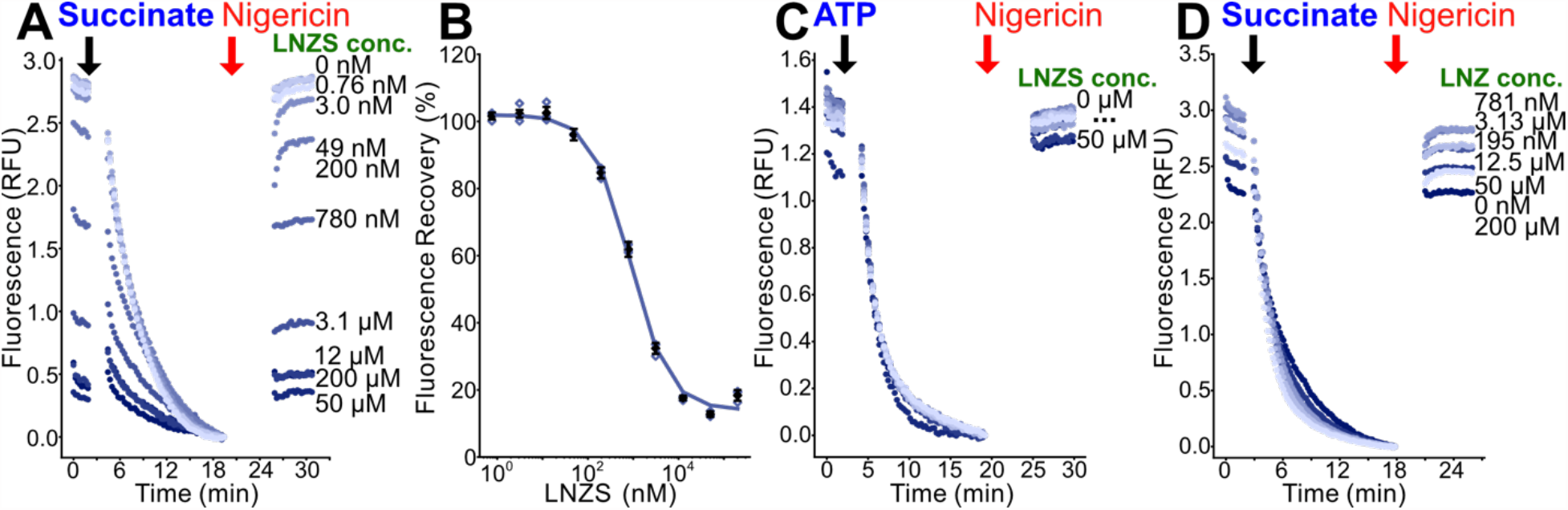
Lansoprazole sulfide inhibits CIII_2_CIV_2_. **A**, Lansoprazole sulfide (LNZS) at different concentrations inhibits succinate-driven acidification of IMVs. **B**, A plot of fluorescence recovery versus inhibitor concentration indicates an IC_50_ of ∼860 nM. **C**, LNZS does not inhibit ATP-driven acidification of IMVs, showing that its effect on succinate-driven acidification is not a result of uncoupling the PMF. **D**, Lansoprazole (LNZ) does not inhibit succinate-driven acidification of IMVs, showing that it is not an inhibitor of CIII_2_CIV_2_. Open symbols show technical replicates. Filled symbols show the mean from n=3 technical replicates. Error bars indicate ± s.d. when shown.

Thioridazine (THZ), a first-generation antipsychotic drug, was proposed to be an inhibitor of NDH-2 (Weinstein et al., 2005). THZ has been used in combination with first-line antibiotics to treat extensively drug resistant TB (Amaral and Viveiros, 2017). We found that THZ inhibited both succinate-driven IMV acidification (Fig. 6A and B) and ATP-driven IMV acidification (Fig. 6C and D). Although this result cannot exclude the possibility that THZ inhibits NDH-2, it shows that the present assay is not capable of distinguishing NDH-2 inhibition by THZ from non-specific uncoupling of the PMF. THZ’s micromolar uncoupling activity is consistent with reports that it uncouples oxidative phosphorylation in mitochondria at similar concentration (Rodrigues et al., 2002). Uncoupling of the PMF is an area of active investigation in the study of compounds that sterilize *M. tuberculosis* infection (Hards et al., 2018, 2015; Rao et al., 2008).

**Figure 6.**
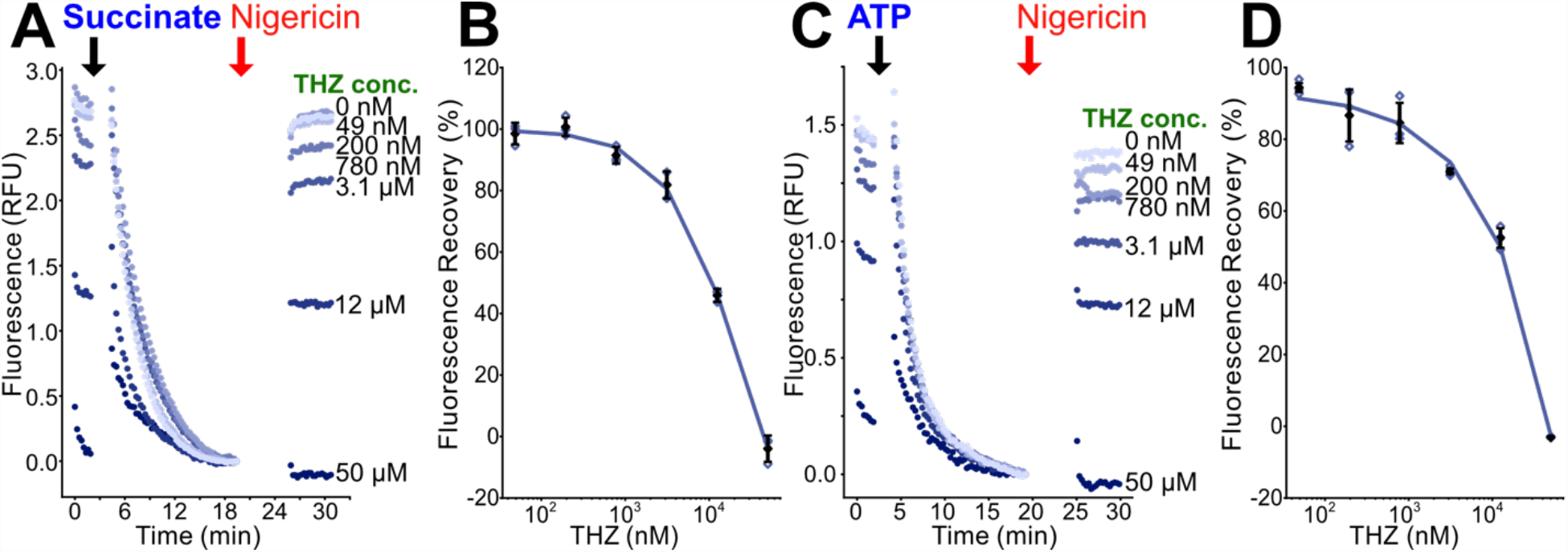
THZ uncouples the PMF in IMVs. **A**, Thioridazine (THZ) at different concentrations inhibits succinate-driven acidification of IMVs. **B**, A plot of fluorescence recovery versus inhibitor concentration for succinate-driven acidification. **C**, THZ at different concentrations inhibits ATP-driven acidification of IMVs, indicating that its effect on succinate-driven acidification results from uncoupling the PMF. **D**, A plot of fluorescence recovery versus inhibitor concentration for ATP-driven acidification. Open symbols show technical replicates. Filled symbols show the mean from n=3 technical replicates. Error bars indicate ± s.d. when shown.

## Discussion

As described above, micromolar concentrations the CIII_2_CIV_2_ inhibitor telacebec appear to block succinate-driven IMV acidification completely. This finding is somewhat surprising because the cytochrome *bd* oxidase should also contribute to acidification, albeit with fewer protons translocated per electron. One explanation for the full inhibition by telacebec is that there is insufficient cytochrome *bd* activity in the IMVs for it to contribute detectably to acidification. This explanation would suggest that in the *M. smegmatis* growth conditions we used cytochrome *bd* makes only a minor contribution to the PMF. However, this conclusion is inconsistent with our finding that telacebec inhibits NADH-driven acidification only slightly. An alternative explanation for these findings is that there could be some form of channeling that occurs in the mycobacterial electron transport chain: with electrons from succinate dehydrogenase preferentially passing to CIII_2_CIV_2_ and electrons from NDH-2 preferentially passing to cytochrome *bd*. Regardless of the cause of this inconsistency, the inability to detect CIII_2_CIV_2_ inhibition with NADH-driven acidification leads to a limitation of the assay. Comparing NADH- and succinate-driven acidification would allow distinguishing between succinate dehydrogenase, NDH-2, and CIII_2_CIV_2_ inhibitors. Without this ability, the assay cannot distinguish succinate dehydrogenase inhibitors from CIII_2_CIV_2_ inhibitors. It is not clear if the assay can be used to detect NDH-2 inhibitors because the proposed NDH-2 inhibitor that we tested uncoupled the PMF. Further, the intrinsic fluorescence of some compounds may interfere with ACMA fluorescence. For these compounds it may be possible to use an alternative fluorophore, such as acridine orange (Courbon et al., 2023).

Despite these limitations, the assay presented here could have utility in characterizing CIII_2_CIV_2_ inhibitors and ATP synthase inhibitors, or even for performing high-throughput screens. The success of the ATP synthase inhibitor bedaquiline supports the value of targeting it, while other studies suggest limitations in targeting CIII_2_CIV_2_ (Arora et al., 2014; Beites et al., 2019). Compared to existing ATP synthase assays, the assay described here is inexpensive and easy to perform, facilitating its use in large scale studies. There are many remaining fundamental questions about mycobacterial oxidative phosphorylation and how it adapts to different growth conditions (Berney and Cook, 2010; Cook et al., 2014; Harold et al., 2022). The assay presented here could serve as a tool to study these aspects of mycobacterial biology.

## Methods

### M. smegmatis strains and growth

For ATP-driven acidification assays, IMVs were prepared from *M. smegmatis* strain GMC_MSM2 (Guo et al., 2021) where a 3×FLAG affinity tag truncates the α subunits of the ATP synthase following residue Ser518. For succinate-, NADH-, malate-, and fumarate-driven acidification assays, IMVs were prepared from *M. smegmatis* strain QcrB-3×FLAG (Yanofsky et al., 2021), which is identical to GMC_MSM2 except that the ATP synthase α subunits are intact and the 3×FLAG tag is at the C terminus of the QcrB subunit of the CIII_2_CIV_2_ supercomplex. *M. smegmatis* strains were grown in Middlebrook 7H9 broth (4.7 g/L) supplemented with 10 g/L tryptone, 2 g/L glucose, 0.8 g/L NaCl, and 0.5 % (v/v) Tween-80. Each 1 L culture was grown in a 2.8 L Fernbach flask at 30 °C for 72 h with shaking at 180 rpm. Bacteria were harvested by centrifugation at for 20 min at 6500 ×g and 4 °C and were sometimes frozen at -80 °C before use.

### Preparation of IMVs

To prepare IMVs from *M. smegmatis* strain QcrB-3×FLAG, 4 L cultures were grown. Cell pellets were resuspended with a Dounce homogenizer in ∼40 mL lysis buffer (50 mM Tris-HCl pH 7.5, 150 mM NaCl, 5 mM MgSO_4_, 5 mM benzamidine hydrochloride, 5 mM 9-aminocaprioc acid) per 1 L of starting cell culture. DNase I in water and PMSF in ethanol were added to the suspension to final concentrations of 100 µg/mL and 100 µM from 100 mg/mL and 100 mM stocks, respectively. The cell suspension was then filtered through four layers of Miracloth (Millipore) to remove clumps of cells and cells were lysed with four passes through an Avestin homogenizer operating at 20,000-25,000 psi. Insoluble debris was removed by centrifugation for 30 min at 39,000 ×g and 4 °C. The membrane fraction from cells was collected by centrifugation for 1 h at approximately 199,269 ×g and 4 °C using a Beckmann Ti70 ultracentrifuge rotor. To form IMVs, pelleted membranes were resuspended with a Dounce homogenizer in 2.5 mL resuspension buffer A (50 mM Tris pH 7.5, 150 mM NaCl, 5 mM MgSO_4_, 5 mM benzamidine hydrochloride, 5 mM 9-aminocaprioc acid, 20 % [v/v] glycerol) per 1 L of original cell culture. IMVs were divided into aliquots and stored at -80 °C. These IMVs were diluted before use, as described below. IMVs from *M. smegmatis* strain GMC_MSM2 were prepared in the same way as from the QcrB-3×FLAG strain with the minor modification that membranes were resuspended at 2.7 mL/L of starting culture in resuspension buffer B (20 mM HEPES-KOH pH 7.5, 50 mM NaCl, 50 mM KCl, 5 mM MgSO4, 20 % [v/v] glycerol).

### Vesicle acidification assays

For succinate-driven acidification experiments with inhibitors, IMVs from the QcrB-3×FLAG strain were diluted 2-fold in resuspension buffer A before use. For NADH-driven acidification experiments with telacebec, IMVs were diluted 32-fold in resuspension buffer A before use. For ATP-driven acidification experiments with inhibitors, IMVs from the GMC_MSM2 strain were used without further dilution. IMV acidification experiments were performed in 96-well plates (BRANDplates™ pureGrade 96-well black microplates). For each well of the plate, 120 µL ACMA assay buffer (20 mM HEPES-KOH pH 7.5, 200 mM KCl, 10 mM MgSO_4_), 35 µL of MilliQ water, and 4.8 µL of inhibitor in DMSO at 50× its intended final concentration, or DMSO alone, was mixed in a microcentrifuge tube at room temperature. To this solution, 0.375 µL of 9-amino-6-chloro-2-methoxyacridine (ACMA) from a 2 mM stock in ethanol and 15 µL of diluted IMV solution were added. Each sample was mixed by pipetting, and 116.8 µL was transferred to each well of the 96 well plate and incubated in the dark for ∼15 min at room temperature prior to starting the experiment.

ACMA fluorescence was followed with a Neo2 Multi-Mode Assay Microplate reader (Synergy) with samples held at 25 °C during experiments. The samples were excited at 410 nm and fluorescence was measured at 480 nm with the fluorescence gain set to 90. Fluorescence of the samples was monitored for 2 min to determine a baseline, at which point 40 µL of either 4 mM disodium NADH in water, 20 mM sodium succinate in water, 20 mM fumarate in 40 mM Tris with pH unadjusted (resulting in a pH of ∼7), 20 mM malate in 40 mM Tris-HCl with pH unadjusted, or 8 mM disodium ATP in 16 mM Tris with pH unadjusted (resulting in a pH of ∼7) was added to each well using the instrument’s automated injector, which had been primed with these solutions. Adding these solutions resulted in final concentrations of 1 mM NADH, 5 mM succinate, 5 mM fumarate, 5 mM malate, or 2 mM ATP, respectively. Fluorescence from the sample was monitored for 15 min before 3.2 µL of 100 µM nigericin in 1% (v/v) ethanol was added to each well with a multichannel pipettor. The samples were then mixed with a separate multichannel pipettor and recovery of fluorescence signal was monitored for an additional 5 min before ending the experiment. The difference between the fluorescence immediately before and 5 min after addition of nigericin is reported as the fluorescence recovery. For visualization, raw fluorescence values were normalized by addition of an offset so that the fluorescence immediately prior to nigericin injection is set to 0.

## Acknowledgments

SAH was supported by a SSuRe scholarship from the Hospital for Sick Children, GMC was supported by a Mary H. Beatty Fellowship from the Department of Medical Biophysics, YL was supported by a Doctoral Canada Graduate Scholarship from the Canadian Institutes of Health Research, and JLR was supported by the Canada Research Chairs program. This work was supported by Canadian Institutes of Health Research Project Grant PJT162186.

## Author contributions

SAH optimized and performed the assays and analyzed the results. GMC and YL taught SAH how to culture *M. smegmatis*, prepare IMVs, and perform assays based on experiments developed previously by GMC. JLR conceived the project. GMC, YL, and JLR supervised the research. SAH and JLR wrote the manuscript and prepared the figures with input from the other authors.

## The authors declare no competing interests

## Figures

**Supplementary Figure 1.**
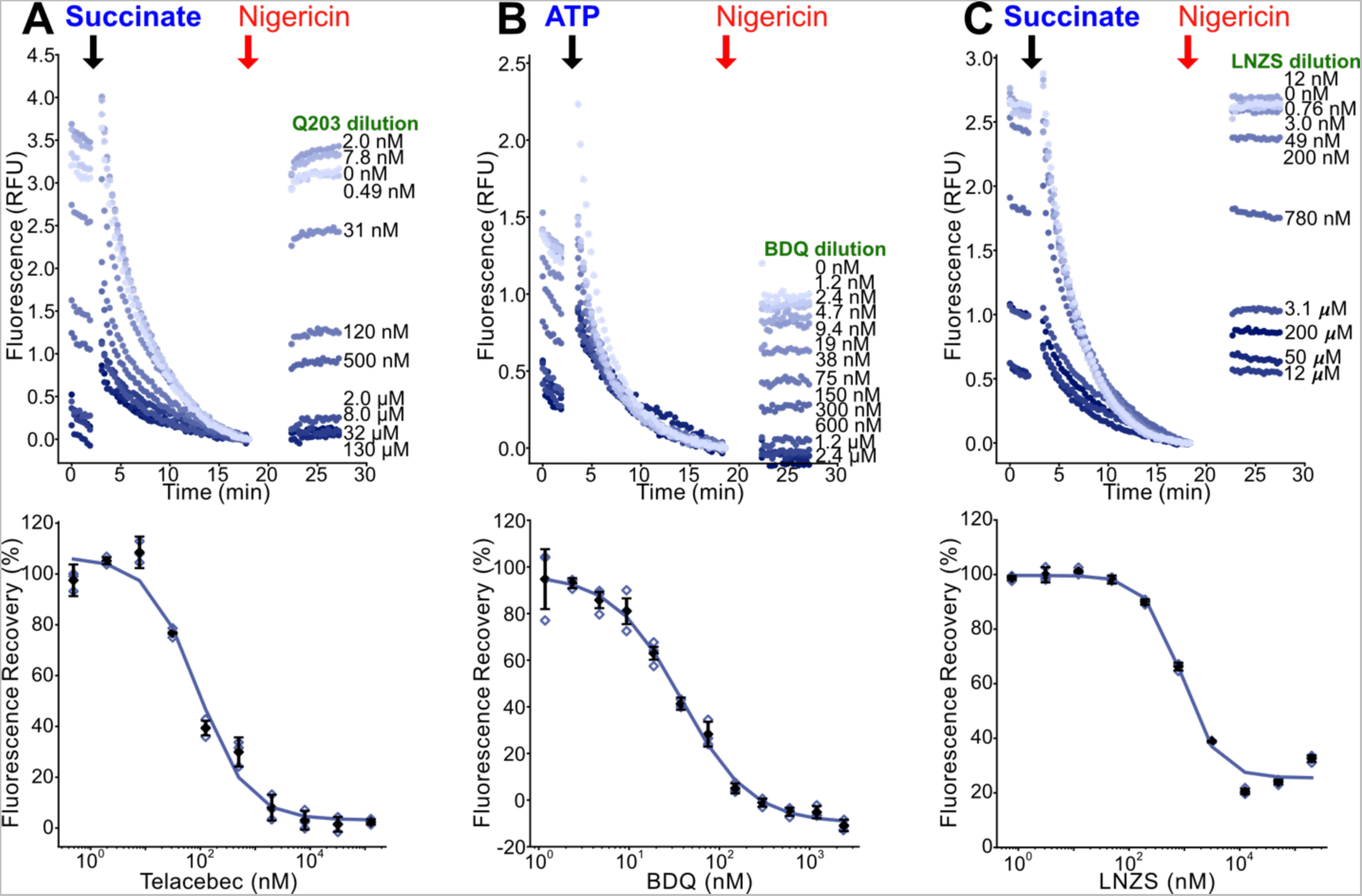
Replication of inhibitor dilution experiments. **A**, Replicate experiment for telacebec (Q203) inhibition of succinate-driven IMV acidification using an independently prepared dilution series of telacebec. The experiments indicate an IC_50_ of ∼90 nM. **B**, Replicate experiments for bedaquiline (BDQ) inhibition of ATP-driven IMV acidification using an independently prepared dilution series of bedaquiline. The experiments indicate an IC_50_ of ∼40 nM. **C**, Replicate experiments for lansoprazole sulfide (LNZS) inhibition of succinate-driven IMV acidification using an independently prepared dilution series of LNZS. The experiments indicate an IC_50_ of ∼890 nM. Open symbols show technical replicates. Open symbols show technical replicates. Filled symbols show the mean from n=3 technical replicates. Error bars indicate ± s.d. when shown.

